# Supplemental irrigation during heat waves affects yield but not whole-vine carbohydrates in wine grapes

**DOI:** 10.64898/2026.06.24.734398

**Authors:** Morgan E. Furze, Alan Rodriguez-Urquidi, Martina Galeano, Joseph Lozano, Luis Sanchez, Nick Dookozlian, Andrew J. McElrone, Elisabeth J. Forrestel

**Affiliations:** Department of Botany and Plant Pathology, Purdue University, West Lafayette, IN 47907; Department of Forestry and Natural Resources, Purdue University, West Lafayette, IN 47907; Center for Plant Biology, Purdue University, West Lafayette, IN 47907; Department of Viticulture and Enology, University of California, Davis, USA 95616; Winegrowing Research, E&J Gallo Winery, Modesto CA USA 95354; USDA-Agricultural Research Service, Davis, CA USA 95616

**Keywords:** heat waves, irrigation, nonstructural carbohydrates, perennial crops, resilience, wine grapes, yield

## Abstract

As extreme heat events increase in frequency and intensity worldwide, understanding how woody perennial crops respond to higher maximum temperatures is critical. Perennials face distinct challenges, persisting across many seasons under increasingly variable and extreme conditions, and heat waves threaten the viability of wine grape cultivars through impacts on yield, wine quality, and long-term vine health. To test whether irrigation practices before and during heat waves affect grapevine carbon (C) storage and health, we experimentally manipulated irrigation regimes surrounding heat waves from 2019–2021 in a commercial Cabernet Sauvignon vineyard in the Lodi AVA of California’s Central Valley. Vine physiological traits and yield were measured throughout, and whole-vine nonstructural carbohydrate (NSC) concentrations were quantified after three growing seasons. Although lower supplemental irrigation reduced photosynthesis, stomatal conductance, and fruit yield, whole-vine NSCs did not differ significantly in any perennial organ by the experiment’s end, indicating that reproductive output and final NSC status responded to irrigation on different timescales. These results suggest that moderate supplemental irrigation during heat events is sufficient to mitigate negative impacts on yield and quality while supporting recovery of NSC reserves, though longer-term monitoring is needed to confirm that this short-term resilience persists.

## Introduction

Extreme heat and drought conditions have increased globally in recent decades ((Mukherjee and Mishra 2021; Mazdiyasni and AghaKouchak 2015). These events, often intrinsically linked with each other, pose a significant threat to plants in natural and managed ecosystems (Wu and Jiang 2022; Ruffault et al. 2018; Zampieri et al. 2017; Allen et al. 2010). In agroecosystems, crops have exhibited a diversity of responses to heat wave (HW) and drought events including stunted growth, altered phenology, and mortality (Hatfield and Prueger 2015; Parker et al. 2024), with repercussions for reduced quality and yield that may lead to economic losses and food insecurity (Reyes and Elias 2019; Campbell et al. 2026; Galeano et al. 2026). For woody perennial crops in particular, their capacity to recover following stress determines their persistence and function over time, and they may have a resilience or recovery advantage over those in natural systems due to management tools such as irrigation. However, we have a limited understanding of how the timing and amount of irrigation impacts recovery dynamics following extreme heat as well as long-term perennial crop health.

In arid climates, where high quality grapes are most commonly grown, irrigation is used as a management tool to maintain quality and sustain crop water demand during warm, dry summers. Some perennial crops such as grapevines are well-adapted to semi-arid conditions, and controlled or regulated deficit irrigation (RDI) can aid in managing vine vigor, enhancing berry quality and yield, while also reducing water use in water-scarce areas like California and other Mediterranean climates (Kuhn et al. 2014; Chaves et al. 2010, 2007). Nevertheless, extreme heat exposure exacerbates water stress, causing stomatal closure and reduced carbon (C) assimilation, potentially leading to a tipping point in which source-sink imbalances and other physiological dysfunction (e.g. hydraulic failure) slow or even prevent vine recovery (Prats et al. 2023; Lopez-Fornieles et al. 2022). Previous studies have shown that pre-veraison and mid-ripening heat exposure decreased berry size and fresh weight in grapevines, and further affected the dynamics of primary and secondary metabolites, such as sugars and flavonoids, which ultimately shape wine quality (Buttrose et al. 1971; Greer and Weston 2010; Kuhn et al. 2014; Gouot et al. 2019; Yu et al. 2020; Gambetta et al. 2020; Campbell et al. 2026; Galeano et al. 2026). Such immediate physiological changes may be alleviated by adapting the amount and timing of irrigation to replace adequate crop evapotranspiration (ET) demands during HWs, but carryover effects on long-term vine physiology and health deserve further attention.

Nonstructural carbohydrates (NSCs) have been commonly used as an indicator of crop resilience and health (e.g., (Zwieniecki et al. 2022; Purdy et al. 2015; Gorlitsky et al. 2015). NSCs, mainly sugars and starch, serve as substrates for critical processes like growth, osmoregulation, and metabolism (Hartmann and Trumbore 2016). Previously stored NSCs can be remobilized when demand exceeds supply, thus, NSC status plays an important role in survival and recovery from stress. In leaves, low NSC levels may result from reduced photosynthetic rates following heat and water stress (Tombesi et al. 2021; Chaves et al. 2002; Maroco et al. 2002). NSCs have also been found to decline in the petioles and trunks of water-stressed wine grapes, possibly signaling C limitation (Vuerich et al. 2023; Falchi et al. 2020). In contrast, high soluble sugar levels may accumulate to maintain cell turgor under heat and drought stress and to facilitate recovery (Long and Adams 2023; Klein et al. 2018; Chitarra et al. 2014; Chen and Murata 2002; Brodersen and McElrone 2013). Overall, shifts in whole-vine C partitioning to biomass and reserves during the growing season have been documented in response to water deficit and soil temperature (Torres et al. 2021; Vuerich et al. 2021; Field et al. 2020), yet the long-term impact that abiotic stress, especially extreme heat exposure, has on whole-vine C balance is limited. Without empirical evidence elucidating the long-term physiological consequences of HWs, our ability to select resilient varieties and improve irrigated agricultural systems under future climates will be hindered, and thus there exists an opportunity to ask the critical question – can we minimize water use surrounding HWs without detrimental effects on vine health as indicated by NSC status, yield, and berry quality?

To determine whether different irrigation practices surrounding HWs impact the long-term C allocation and health of grapevines, we manipulated irrigation during HW events during the 2019, 2020, and 2021 growing seasons using a variable rate drip irrigation system in a commercial vineyard (Lodi, CA, USA) planted with *Vitis vinifera* cv. Cabernet Sauvignon, the world’s most commonly grown red grape variety. Three supplemental irrigation treatments—initiated one to two days prior through the end of the HW—were implemented: baseline (deficit irrigation, 60% ET), 1.5-2x baseline (90-120% ET), and 2-3x baseline (120-180%). At the end of three growing seasons, we measured aboveground and belowground NSC concentrations to assess whether long-term NSC status was impacted by supplemental irrigation treatments. Due to the expected negative impacts on vine physiology (e.g., reduced C assimilation) following multiple HW events, we hypothesized that there would be long-term impacts on NSCs after three growing seasons, such that NSC concentrations would be lower for vines in the baseline compared to those receiving supplemental irrigation (i.e., baseline < 2x current ET < 3x current ET).

## Materials and Methods

### Field site

This study was carried out from 2019 through the 2021 growing seasons in a commercial vineyard located in Lodi, CA, USA (latitude: 38° 17’ 37.1” N, longitude: 121° 06’ 39.7” W, elevation: 40-50 masl), one of the most productive grape growing regions of the Central Valley. The site is a 23-acre block of 9-year-old *Vitis vinifera* cv. Cabernet Sauvignon grafted on 1103 Paulsen as the rootstock. Rows are oriented East-West with a vine spacing of 1.8 m and a row spacing of 3 m.

The vineyard block was divided into a total of 98 subplots of 30 m x 30 m with a variable rate drip irrigation (VRDI) system allowing the application of a differential irrigation to each individual subplot (Sanchez et al. 2017). Applied irrigation was scheduled throughout the growing season using a method that calculates crop evapotranspiration (ET_c_) using a combination of both ET_o_ from a nearby weather station and a dynamic, modified crop coefficient (K_c_) approach (Allen et al. 1998). K_c_ values were derived by using remotely-sensed normalized difference vegetation index (NDVI) specific to each 30 m x 30 m variable rate zone, along with empirical parameters representing the expected water demands for a non-stressed vineyard.

### Experimental design and treatment application

In the context of this experiment, a HW was identified based on historical weather data and trends as a minimum of three consecutive days with maximum temperatures above 38° C. Recognizing that heat extreme parameters are defined relative to local historical conditions, the maximum temperature cutoff of 38° C was determined using the 1980-2010 climate as the baseline, the 95% percentile of maximum temperatures during the growing season (April - October) were used to determine temperature above which was considered an extreme heat day. Maximum temperatures were recorded using the Borden Ranch vineyard (38° 17’ 24” N 121° 7’ 12” W) weather station from the GRAPEX project (Kustas et al. 2022), which is located 0.8 kilometers from the field experiment.

In 2019, three subplots (30 m x 30 m) per treatment were randomly selected across the vineyard. In 2020 and 2021, each replicate was expanded to a 60 m x 60 m plot (four 30 m x 30 m subplots). ET was estimated throughout the season using Landsat data, normalized difference vegetation index (NDVI) and crop coefficient (Sanchez et al. 2017). Total irrigation applied throughout the study period varied across the treatments (Fig. 1). Differential irrigation treatments were applied only when a heat event took place and started one or two days before each HW and continued through the last day of the heat event. Three irrigation treatments were implemented during 2019 and 2020: a control or baseline, which was exposed to deficit irrigation and held at 60% ET, a second treatment where the irrigation was double the baseline (2x baseline ET), and a third treatment with triple the amount of water of the baseline (3x baseline ET). In 2021, irrigation treatments were adjusted to 60% ET, 90% ET, and 120% ET during HWs based on the growers’ realization, based on our findings, that excessive water was being used in the 3x baseline ET treatment.

**Figure 1.**
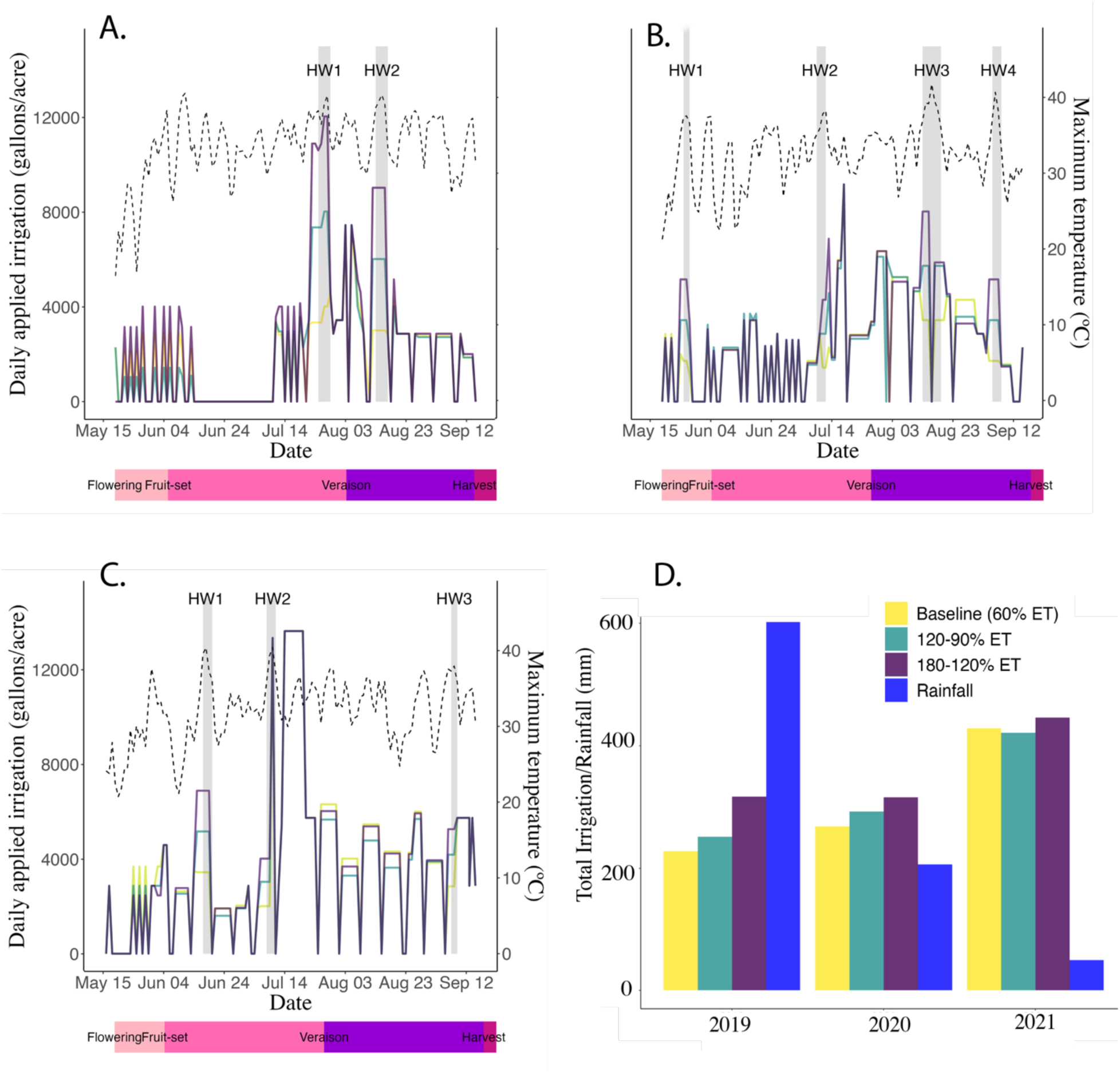
Daily applied irrigation (left axis) for each of three treatments and maximum daily temperature (right axis) during the (A) 2019, (B), 2020, and (C) 2021 growing seasons. Heatwave events (3 or more days predicted to have a maximum temperature above the 38 degrees Celsius threshold) are depicted as gray vertical bars and labeled HW#. Total cumulative irrigation applied and precipitation totals annually for 2019, 2020, and 2021 are displayed in panel (D).

### Canopy NDVI

Canopy reflectance was acquired by aerial multispectral imaging (Ceres Imaging, Oakland, CA, USA) from fixed-wing overflights of the vineyard on dates spanning the (May through mid-September) growing seasons of 2019 (9 flights), 2020 (8 flights), and 2021 (9 flights). For each flight, the normalized difference vegetation index (NDVI) was computed from the provider’s calibrated near-infrared and red reflectance bands as NDVI = (NIR – Red)/(NIR + Red), following Ceres Imaging’s standard radiometric and geometric processing and atmospheric correction. The native imagery (ground sample distance ≈ 3 m) was aggregated to a 30 m × 30 m grid aligned to the experimental layout, and a single mean NDVI value was extracted for each 30 m × 30 m pixel on each date by averaging all finer-resolution pixels falling within it. Twenty-eight pixels overlying the irrigation-treatment plots were retained for analysis (n = 10, 9, and 9 for the 60%, 120%/90%, and 180%/120% ET treatments, respectively), each pixel corresponding to one block × sub-block experimental unit, yielding one NDVI value per treatment pixel per date.

### Yield data collection

Grapes were considered to have reached maturity when total soluble solids (TSS) values were 25 ± 1° Brix. For 2019, six vines per subplot were hand-picked and harvested to estimate yield. Total number of bunches and total fruit fresh weight per vine was recorded. A total of three subplots per treatment were harvested. For 2020 and 2021, twenty-seven vines per subplot were harvested with a total of three subplots per treatment. Berry weights were measured for samples aggregated across vines within sample plots and divided into 60 berry samples for subsequent chemical analysis (see Campbell et al. 2026).

### Sample collection and NSC analysis

Samples were collected from vines (n=54) immediately following harvest over the course of two days in late-September 2021. Leaves, arms, canes, a trunk/stem core, coarse roots, and fine roots were collected from each vine. Two sun leaves, two arms, and two canes were collected (one from each horizontal cordon) and combined into single samples for NSC analysis. Using hand pruners, we collected a 5 cm length of cane between the 1st and 2nd nodes and a 2 cm length of arm directly beneath and kept the bark on both. To obtain the trunk/stem core, we used a standard 4.3-mm increment borer (Haglöf Company Group, Långsele, Sweden) on the south face of each vine and saved the outer 2 cm for analysis. Coarse and fine roots were accessed by carefully digging around the base of the trunk/stem with a shovel. All samples were kept on dry ice in the field and then stored at -80°C.

To prepare samples for NSC analysis, samples were placed in a drying oven at 100°C for 1 hour to deactivate starch degrading enzymes, and then at 70°C for 2–3 days until the samples were completely dry. Dried samples were then ground (mesh 20; Thomas Scientific Wiley Mill, Swedesboro, NJ, USA) and 30 mg of each homogenized sample was weighed into a 2-ml microcentrifuge tube for sugar and starch analyses.

For sugar and starch analyses, we followed the routine procedures detailed in (Landhäusser et al. 2018). For sugar analysis, samples were extracted with ethanol in a 90°C water bath and the resulting sugar extracts were subjected to a phenol-sulfuric acid reaction and measured at 490 nm on a spectrophotometer (Agilent TechnologiesCary 100 UV-Vis). For starch analysis, the tissue pellet was left to dry under the fume hood overnight following sugar extraction and was then sequentially digested with α-amylase and amyloglucosidase to produce glucose hydrolysate. Peroxidase-glucose oxidase color reagent and 75% sulfuric acid were then used in the reaction and the resulting solutions were measured at 525 nm on the spectrophotometer. Sugar and starch concentrations are expressed as mg per g dry wood and provided in Table S1.

### Statistical analyses

All statistical analyses were performed using R software (Version 4.3.2). To determine if NSC concentrations differed between irrigation treatments, we used a one-way analysis of variance (ANOVA) to analyze sugar concentrations among irrigation treatments for each organ independently. The analysis was repeated for starch concentrations. The same model was used for all analyses and accounted for the number of years in a treatment, a blocking factor, and replicates within a block. Differences between pairs of means were evaluated with Tukey’s honest significant difference (HSD) at α = 0.05. Similarly, to compare yield components between irrigation treatments, we used one-way ANOVAs for yield, cluster weight, cluster counts, and berry weight independently and Tukey’s HSD at α = 0.05. NDVI differences in irrigation treatments were evaluated for each date of sampling independently using one-way ANOVAs.

## Results

### Heat waves and treatment applications during the growing seasons

Across the three growing seasons (2019-2021), a total of nine HWs were recorded (Fig. 1A-C). The 2019 growing season had two HWs, one at the end of July and another in mid-August. The 2020 growing season had four HWs at the end of May, mid-July, mid-August, and early September. Three HWs occurred during the 2021 growing season in mid-June, early July, and early September. Cumulative irrigation increased from 2019-2021 with differences corroborating the irrigation treatments, such that the 3x baseline ET treatment plots had higher cumulative irrigation than the 2x baseline ET treatment and baseline plots (Fig. 1D).

### Canopy NDVI

Canopy NDVI followed the expected phenological trajectory in all three seasons, rising from ∼0.50 in early May to a peak near 0.70–0.80 in June, declining through post-veraison in July and August, and rising again toward harvest in September (Table S1). For all flight dates across all years, NDVI did not differ significantly among the three irrigation treatments (one-way ANOVA across the 28 treatment pixels; all P > 0.05). Treatment means typically fell within 0.01– 0.03 NDVI units of one another—well within the among-pixel standard error—and the three treatment trajectories overlapped throughout each season (Table S1).

### Yield components

In 2019 and 2020, the 2x baseline ET treatment had significantly higher yields than the baseline (Table 1). The same pattern held in 2021—with the lowest yield in the baseline—but was not significant (Tukey baseline vs. 2x, P = 0.06), likely because the contrast was weakened by a narrower irrigation range (90% and 120% ET vs. 120% and 180% ET), similar amounts of total irrigation applied that season (Fig. 1D), and greater plot-to-plot variability. In 2019 and 2020, applied irrigation beyond 2x resulted in no additional yield increase. The lower baseline yield was driven mainly by smaller berries, significant in both 2019 and 2020. Although the treatment difference in cluster weight was significant only in 2019, cluster weight was the largest contributor to the size of the baseline-to-2x yield gap in both years (∼60% in 2019, ∼70% in 2020), even though the between-treatment difference fell short of significance in 2020. Cluster number per plant never differed significantly, though in 2021 it tended to rise with supplemental irrigation (lowest in the baseline, highest under 2x).

**Table 1.** Yield components for the 2019-2021 growing seasons taken at harvest of vines exposed to different irrigation treatments prior to and during HWs. Error bars denote 1 ± SE of the mean. Note: 2x baseline ET and 3x baseline ET for 2019-2020 and 1.5x baseline ET and 2x baseline ET for 2021 as detailed in the Materials and Methods.

| 2019 |  |  |  |  |
| --- | --- | --- | --- | --- |
|  | Baseline (60% ET) | 2x Baseline ET | 3x Baseline ET | P-value |
| Yield (kg/plant) | $6.33 \pm 0.21^b$ | $8.86 \pm 0.63^a$ | $7.50 \pm 0.49^{ab}$ | 0.007 ** |
| Clusters (no./plant) | $94.2 \pm 3.6^a$ | $105.7 \pm 4.6^a$ | $95.5 \pm 5.1^a$ | 0.196 ns |
| Weight (gr/cluster) | $67.99 \pm 2.61^b$ | $83.53 \pm 4.63^a$ | $79.09 \pm 3.40^{ab}$ | 0.028 * |
| Berry weight (gr) | $0.869 \pm 0.002^b$ | $1.046 \pm 0.012^a$ | $1.027 \pm 0.003^a$ | <0.001 *** |
| 2020 |  |  |  |  |
|  | Baseline (60% ET) | 2x Baseline ET | 3x Baseline ET | P-value |
| Yield (kg/plant) | $7.22 \pm 0.15^b$ | $8.28 \pm 0.30^a$ | $8.18 \pm 0.34^{ab}$ | 0.027 * |
| Clusters (no./plant) | $98.4 \pm 2.4^a$ | $102.2 \pm 2.2^a$ | $103.2 \pm 3.7^a$ | 0.418 ns |
| Weight (gr/cluster) | $75.09 \pm 2.54^a$ | $82.27 \pm 2.94^a$ | $80.35 \pm 2.02^a$ | 0.106 ns |
| Berry weight (gr) | $0.966 \pm 0.005^b$ | $1.011 \pm 0.007^a$ | $1.002 \pm 0.012^a$ | <0.001 *** |
| 2021 |  |  |  |  |
|  | Baseline (60% ET) | 1.5x Baseline ET | 2x Baseline ET | P-value |
| Yield (kg/plant) | $7.35 \pm 0.54^a$ | $8.89 \pm 0.54^a$ | $8.26 \pm 0.66^a$ | 0.158 ns |
| Clusters (no./plant) | $96.8 \pm 6.2^a$ | $106.3 \pm 4.0^a$ | $110.3 \pm 3.0^a$ | 0.102 ns |
| Weight (gr/cluster) | $80.43 \pm 7.94^a$ | $85.72 \pm 4.83^a$ | $75.97 \pm 5.48^a$ | 0.514 ns |
| Berry weight (gr) | $1.087 \pm 0.014^a$ | $1.100 \pm 0.008^a$ | $1.057 \pm 0.015^a$ | 0.052 ns |

Values are mean ± SE. P-values are from one-way ANOVA (2019) or a likelihood-ratio test of the mixed model (2020–2021); \*, \*\* and \*\*\* denote $P < 0.05$ , $< 0.01$ and $< 0.001$ ; ns, not significant. Different letters denote significant differences among treatments for each variable (Tukey HSD, $P < 0.05$ ).

### Organ-level nonstructural carbohydrate concentrations

Despite differential irrigation prior to and surrounding HWs, grapevine NSC concentrations, which were assessed in September 2021 after three growing seasons, did not differ between the irrigation treatments. There were no differences in sugar or starch concentrations across the irrigation treatments for any organ assessed, including leaves, canes, arms, trunks, coarse roots, and fine roots (Fig. 2).

**Figure 2.**
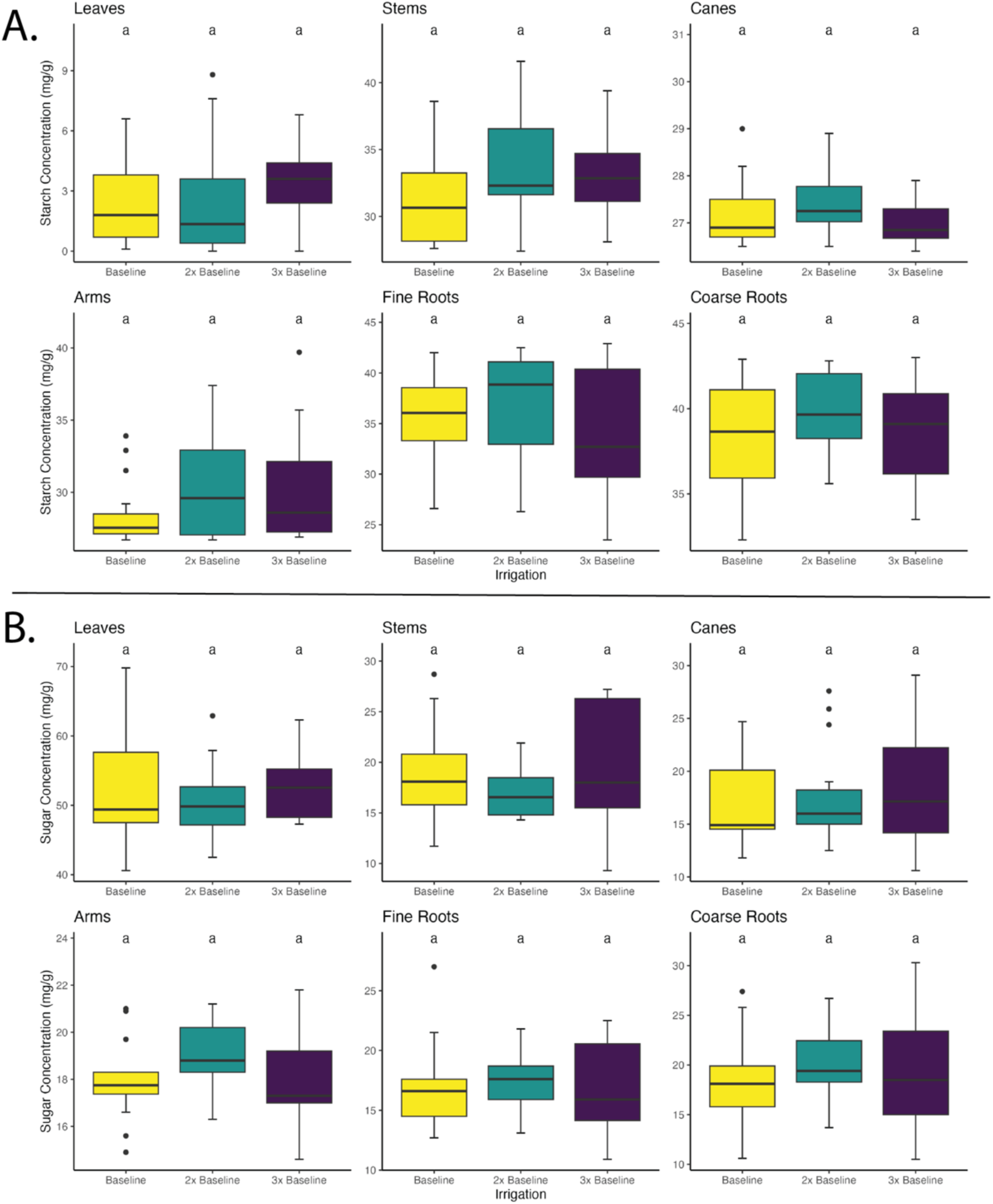
Sugar (A) and starch (B) concentrations (mg/g) in grapevine organs following three irrigation treatments prior to and surrounding HWs throughout 2019-2021. Lowercase letters indicate significance of differences among irrigation treatments. Trunk and stem are used interchangeably.

Further, as previously detailed in the *Materials and Methods* section, each replicate was expanded from three pixels (30 m x 30 m) per treatment to four pixels, or a 60 m x 60 m plot, in 2020, resulting in an irrigation treatment duration of 3 years (2019-2021) versus 2 years (2020-2021) for some vines. Treatment duration had no impact on organ-level NSC concentrations, with the exception of cane sugar concentrations which were higher in the irrigation treatment duration of three years, as compared to two years (Fig. S1). Overall, our assessment of whole-vine NSC status at the end of the irrigation manipulation experiment indicated that the highest sugar concentrations were observed in the leaves (Fig. 2A) and the highest starch concentrations were observed in the fine and coarse roots (Fig. 2B), but there were no differences across irrigation treatments.

## Discussion

The use and replenishment of NSCs have implications for the long-term health and survival of perennial species. Given the expected negative impacts on vine physiology following repeated HW exposure—chiefly reduced C assimilation—we predicted that lower supplemental irrigation would progressively deplete NSCs, leaving deficit (baseline) vines with lower end-of-season NSCs than vines receiving 2x or 3x baseline ET. This was not the case. Quantifying NSCs across all major vine organs—leaves, canes, arms, trunk, and coarse and fine roots—at the end of three growing seasons, we found that whole-vine NSC status did not differ among irrigation treatments, indicating recovery to a common level by the end of the study regardless of irrigation manipulation.

Complementary measurements from the same vineyard and treatments confirm that the irrigation regimes imposed consistent physiological responses: relative to supplemental irrigation, the baseline reached significantly more negative predawn and midday water potentials, and exhibited reduced net photosynthesis, stomatal conductance, and leaf cooling during and immediately after HWs (Galeano et al. 2026). Because the deficit was imposed only within narrow windows before and during HWs, vines received standard—and in some years more abundant—irrigation through the remainder of each season and after harvest. NSCs therefore had ample opportunity to replenish, and the convergence in end-of-season NSCs is most parsimoniously interpreted as post-stress replenishment that equalized NSCs across treatments, rather than insensitivity to treatments that measurably depressed water status, gas exchange, yield, and berry quality (Campbell et al. 2026; Galeano et al. 2026). Critically, however, this recovery did not eliminate the cost of lower irrigation: yield and berry quality differences persisted—including between treatments receiving the same total seasonal water but differing in its timing—indicating that episodic water limitation impacted reproductive output even when it left no lasting imprint on NSC storage (Table 1, (Campbell et al. 2026). Soluble sugars were highest in the leaves and starch was highest in the coarse and fine roots (Fig. 2), yet no organ differed consistently among treatments. The single exception was a modest elevation of cane sugar under three versus two years of treatment (Fig. S2), pointing toward accumulation rather than depletion under prolonged manipulation (Pichierri et al. 2026). Thus, consistent NSCs across organs likely reflects NSC recovery.

Canopy-scale observations reinforce this interpretation. Remotely sensed NDVI, an integrated proxy for canopy greenness and vegetative health, did not differ significantly among irrigation treatments on any date in any of the three seasons, with the treatment trajectories overlapping throughout each growing season (Table S1). Since canopy greenness was largely maintained even as instantaneous gas exchange declined under deficit (Gal et al. 2025; Galeano et al. 2026), it suggests that transient declines did not register at the whole-canopy scale, consistent with the vines’ subsequent ability to replenish NSC stores despite seasonal impacts on yield and quality.

The cost of episodic deficit thus registered in the fruit rather than in stored C, extending beyond yield to berry composition (Campbell et al. 2026). Sugars act not only as storage and respiratory substrates but as signaling molecules, with both sucrose and glucose altering gene expression in the phenylpropanoid pathway and thereby phenolic production (Wind et al. 2010; Ferri et al. 2011; Solfanelli et al. 2006)—metabolites contributing to pathogen defense, abiotic stress tolerance, and berry and wine quality. The same irrigation treatments that left end-of-season NSCs unchanged nonetheless altered berry quality (Campbell et al. 2026), suggesting that C status may shape fruit composition through regulatory pathways distinct from bulk NSC reserve accumulation, even when whole-vine NSC reserves appear buffered. A vine can thus maintain its NSC reserves while the timing and amount of water around HWs still leave a measurable signature on the harvested crop.

Total yield per vine partitions into cluster number, cluster weight (berries per cluster), and berry weight. Cluster weight contributed most to the proportional gap between baseline and the 2x supplemental irrigation treatment in both 2019 and 2020 (roughly 60% and 70%, respectively), with cluster number making up the rest. In 2019 the cluster-weight share reflected berry size alone: deficit vines produced significantly smaller berries (Table 1). In 2020 it reflected both smaller berries and fewer berries per cluster, though berry weight was again the only component to differ significantly. Berry size was thus the most consistent significant signal across seasons—the current-season, expansion-stage component most directly exposed to water status during and after the heat-wave windows. By 2021 the pattern changed: no component differed significantly, the yield difference was only a non-significant trend—reflecting greater plot-to-plot variability and a narrower irrigation contrast (1.5x/2x versus 2x/3x in 2019-2020)—and the partition shifted from berry size toward cluster number and cluster weight. Cluster number per vine is set largely by inflorescence initiation in the previous season’s buds, a process sensitive to water stress around flowering and a major source of year-to-year yield variation in grapevine (Guilpart et al. 2014). The lower cluster number in baseline vines in 2021 is thus consistent with a potential carry-over effect from the previous seasons.

Overall, the lagged, component-shifting yield response contrasted with the replenishment of whole-vine NSCs to a common level. Such buffering of NSC status is consistent with emerging evidence that the non-metabolic roles of soluble sugars in grapevine are not simple functions of heat and water status (Long and Adams 2023). Sugars contribute substantially to daily osmotic adjustment and turgor maintenance under declining leaf water potential, yet their osmotic contribution declines seasonally and does not scale with imposed water deficit (Perry et al. 2026); likewise, the rate of seasonal osmotic adjustment in vine leaves appears largely independent of ambient temperature and mild water deficit (Farolfi et al. 2025). The fact that the sugar dynamics most plausibly linked to stress tolerance are themselves only weakly coupled to temperature and water status offers a leaf- and cell-scale parallel to our whole-vine observation: bulk NSC reserves do not always track the stressors that measurably affect physiology and yield. Whether this reflects preferential allocation to NSC reserves or simply the ample post-stress replenishment afforded by managed irrigation cannot be resolved here; either way, NSC status is a comparatively buffered dimension of vine response that recovered while the negative effect of deficit on reproduction persisted and shifted across yield components from season to season.

This decoupling of short-term physiological stress from long-term reserve status is consistent with observations in natural systems, where the effect of water limitation on NSCs depends strongly on the stress intensity and duration (Ruehr et al. 2019). Stemwood NSCs in temperate deciduous trees were depleted following the 2016 drought in the northeastern United States, but rebounded quickly in subsequent years (Blumstein and Furze 2022). In a field-based drought experiment on mature piñon, a one-year drought did not affect NSC reserves despite significant reductions in water potential, photosynthetic rates, and growth, whereas a long-term drought (>10 years) reduced sapwood starch concentrations (Peltier et al. 2023). Because stressed plants may shift metabolic substrates over time—from recently fixed C to older reserves and potentially lipids—measurable reserve decline can lag stress onset by years (Helm et al. 2024). Unlike the self-supporting trees in these studies, vines invest little in mechanical support and allocate proportionally more C to leaves, roots, and storage (Putz and Mooney 1991), a C economy that may favor the reserve buffering and rapid replenishment we observed. Recovering reserves across three seasons of repeated heat and water stress suggests our vines remained well short of the source–sink imbalances or potential C-limitation thresholds that slow recovery in more severely or chronically stressed systems. Whether this resilience persists indefinitely, and under variable circumstances, is less certain: if NSCs decline cumulatively and slowly over time, there are implications for whole-vine C trade-offs and their vulnerability to compounding stressors.

We quantified NSCs at a single post-harvest time point, capturing net status entering dormancy rather than the within-season or event-scale dynamics that unfold during and immediately after individual HWs; finer temporal sampling would reveal whether NSCs are transiently drawn down and refilled within a season. The three-year duration may also have been too short to elicit a C-starvation response, and longer studies are needed to assess these trade-offs in agricultural systems and to determine whether vines that appear resilient in the short term accrue costs that surface years later (He et al. 2020; Peltier et al. 2023). Finally, because we constrained water only before and during heat events, our design does not address season-long limitation; if NSCs were drawn down gradually over many years of sustained deficit, subsequent irrigation might not replenish them quickly. Resolving these questions will require tracking physiology, yield, and NSCs together over longer periods and across both episodic and season-long water regimes.

Taken together, our results indicate that moderate supplemental irrigation around HWs is sufficient to maintain whole-vine NSC status over three seasons, and that water inputs beyond a moderate level confer little additional benefit for storage or yield. Returning to the question that motivated this work—whether water use around HWs can be reduced without long-term detriment to vine health—our data suggest that, over this timeframe, NSCs are not the dimension most at risk: episodic deficit left reproductive output and fruit composition altered while NSCs recovered. NSC monitoring may therefore offer a practical indicator for guiding more sustainable input use, provided it is paired with measures of yield and quality that capture costs reserves do not. Whether this short-term resilience persists under longer and more frequent heat exposure remains the central question for sustaining vine longevity under future climates.

## Acknowledgements

This work is supported by the Agricultural and Food Research Initiative Post Doctoral Fellowship, project award nos. 2021-67034-35230 and 2021-67034-39050, from the U.S. Department of Agriculture’s National Institute of Food and Agriculture.

Support from E. & J. Gallo and their staff was critical to the success of the field trial, seed funding from USDA Climate Hub, CDFA Specialty Crop Block Grant Award No. 19-0001-013-SF, Department of Viticulture & Enology and College of Environmental and Agricultural Science at UC Davis.

We thank S. Castro Bustamante, I. Wright, P. Tolentino, R. Halsey, K. Elemendorf, and I. Joseph for their assistance with field collection.

