## Supplemental Files for "Supplemental irrigation during heat waves affects yield but not whole-vine carbohydrates in wine grapes"

**Table S1.** Remotely-sensed canopy Normalized Difference Vegetation Index (NDVI) values (mean ± SE) aggregated for each block (or pixel) on dates across the growing season between May and mid-September for all study years (2019-2021). Significant differences between irrigation treatments were assessed using a one-way ANOVA for each sampling data.

| **2019** | | | | |
| --- | --- | --- | --- | --- |
| **Date** | **Baseline (60% ET)** | **2x Baseline ET** | **3x Baseline ET** | **P-value** |
| 05 May | 0.505 ± 0.015 | 0.504 ± 0.008 | 0.504 ± 0.008 | 0.993 ns |
| 30 May | 0.732 ± 0.012 | 0.722 ± 0.018 | 0.726 ± 0.019 | 0.916 ns |
| 14 Jun | 0.792 ± 0.006 | 0.797 ± 0.005 | 0.796 ± 0.006 | 0.802 ns |
| 04 Jul | 0.649 ± 0.014 | 0.662 ± 0.007 | 0.651 ± 0.011 | 0.645 ns |
| 19 Jul | 0.628 ± 0.013 | 0.638 ± 0.008 | 0.624 ± 0.011 | 0.660 ns |
| 03 Aug | 0.623 ± 0.012 | 0.638 ± 0.006 | 0.626 ± 0.011 | 0.530 ns |
| 18 Aug | 0.648 ± 0.011 | 0.660 ± 0.008 | 0.647 ± 0.010 | 0.591 ns |
| 02 Sep | 0.648 ± 0.011 | 0.665 ± 0.006 | 0.652 ± 0.010 | 0.433 ns |
| 12 Sep | 0.688 ± 0.010 | 0.701 ± 0.006 | 0.687 ± 0.008 | 0.464 ns |
| **2020** | | | | |
| **Date** | **Baseline (60% ET)** | **2x Baseline ET** | **3x Baseline ET** | **P-value** |
| 04 May | 0.500 ± 0.009 | 0.486 ± 0.009 | 0.490 ± 0.013 | 0.598 ns |
| 24 May | 0.681 ± 0.011 | 0.654 ± 0.013 | 0.660 ± 0.015 | 0.300 ns |
| 08 Jun | 0.709 ± 0.006 | 0.712 ± 0.008 | 0.706 ± 0.007 | 0.843 ns |
| 23 Jun | 0.658 ± 0.010 | 0.661 ± 0.012 | 0.653 ± 0.013 | 0.886 ns |
| 08 Jul | 0.609 ± 0.007 | 0.620 ± 0.008 | 0.610 ± 0.008 | 0.553 ns |
| 23 Jul | 0.595 ± 0.007 | 0.601 ± 0.008 | 0.595 ± 0.010 | 0.866 ns |
| 07 Aug | 0.612 ± 0.006 | 0.617 ± 0.008 | 0.609 ± 0.007 | 0.735 ns |
| 27 Aug | 0.686 ± 0.005 | 0.702 ± 0.006 | 0.689 ± 0.007 | 0.151 ns |
| **2021** | | | | |
| **Date** | **Baseline (60% ET)** | **2x Baseline ET** | **3x Baseline ET** | **P-value** |
| 04 May | 0.506 ± 0.009 | 0.482 ± 0.007 | 0.483 ± 0.009 | 0.108 ns |
| 19 May | 0.607 ± 0.010 | 0.595 ± 0.011 | 0.585 ± 0.012 | 0.379 ns |
| 03 Jun | 0.674 ± 0.009 | 0.690 ± 0.008 | 0.665 ± 0.014 | 0.274 ns |
| 18 Jun | 0.624 ± 0.014 | 0.635 ± 0.014 | 0.611 ± 0.017 | 0.548 ns |
| 08 Jul | 0.551 ± 0.012 | 0.567 ± 0.010 | 0.541 ± 0.012 | 0.300 ns |
| 23 Jul | 0.548 ± 0.008 | 0.572 ± 0.006 | 0.546 ± 0.013 | 0.123 ns |
| 22 Aug | 0.654 ± 0.006 | 0.670 ± 0.006 | 0.645 ± 0.012 | 0.122 ns |
| 06 Sep | 0.687 ± 0.013 | 0.688 ± 0.015 | 0.666 ± 0.016 | 0.518 ns |
| 11 Sep | 0.745 ± 0.012 | 0.751 ± 0.013 | 0.728 ± 0.014 | 0.451 ns |

**Figure S1.**  Sugar (A) and starch (B) concentrations of organs for vines exposed to an irrigation treatment duration of 2 years versus 3 years. The difference in irrigation treatment duration for specific vines is detailed in the Materials and Methods. Lowercase letters indicate significance of differences among treatment duration for each organ.

(A)

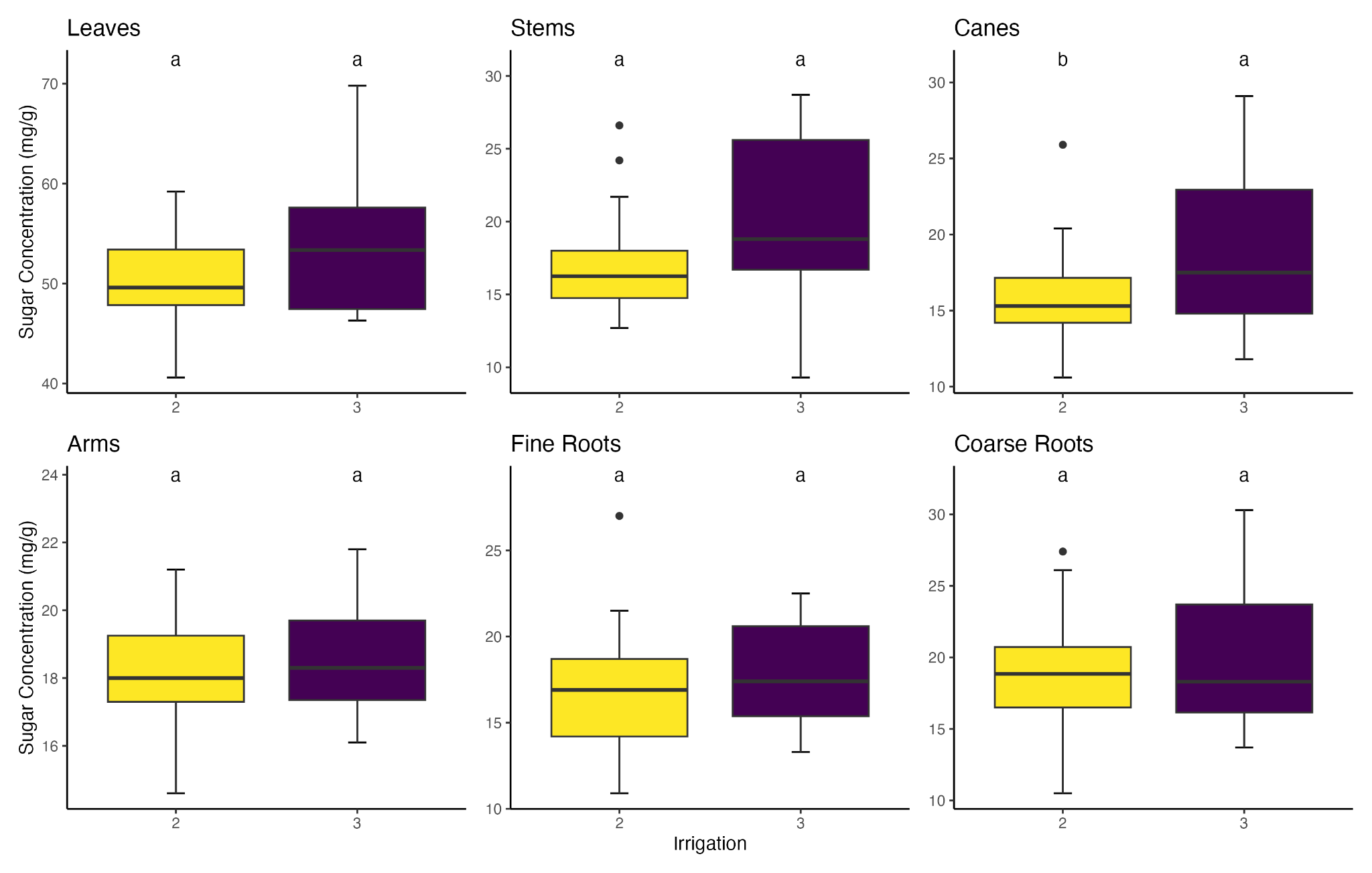

(B)

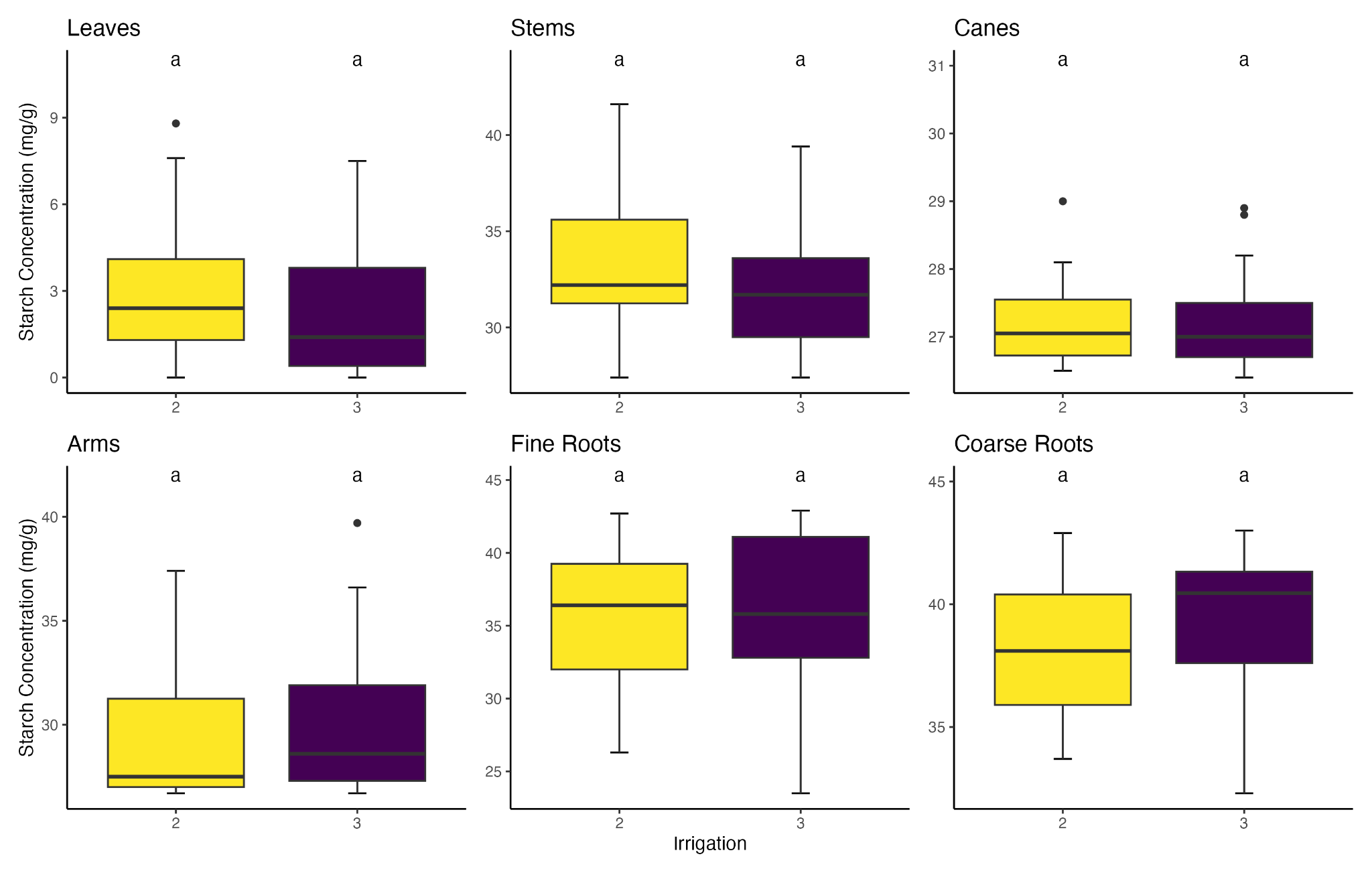

**Table S2.** Mean sugar and starch concentrations in each organ by irrigation treatment and years in the treatment. Treatments are denoted as 60% ET (baseline 60% ET), 120% ET (2x baseline ET), and 180% ET (3x baseline ET). Trunk and stem are used interchangeably.

|  |  |  | Sugar concentration (mg/g) | | Starch concentration (mg/g) | |
| --- | --- | --- | --- | --- | --- | --- |
| Years in treatment | Irrigation treatment | Organ | Mean | SD | Mean | SD |
| 2 | 60% ET | arm | 17.42 | 1.42 | 29.00 | 2.60 |
| 2 | 60% ET | cane | 15.72 | 2.83 | 29.49 | 5.03 |
| 2 | 60% ET | coarse root | 18.67 | 4.72 | 38.17 | 3.61 |
| 2 | 60% ET | fine root | 17.71 | 4.38 | 37.09 | 2.69 |
| 2 | 60% ET | leaf | 50.16 | 6.71 | 2.27 | 2.20 |
| 2 | 60% ET | trunk | 16.96 | 2.90 | 32.93 | 3.62 |
| 2 | 120% ET | arm | 19.33 | 1.36 | 29.51 | 4.09 |
| 2 | 120% ET | cane | 15.33 | 1.93 | 29.08 | 3.79 |
| 2 | 120% ET | coarse root | 20.60 | 3.26 | 35.91 | 6.94 |
| 2 | 120% ET | fine root | 17.40 | 2.35 | 34.73 | 6.00 |
| 2 | 120% ET | leaf | 49.92 | 4.18 | 3.44 | 3.03 |
| 2 | 120% ET | trunk | 18.68 | 7.59 | 34.81 | 4.45 |
| 2 | 180% ET | arm | 17.68 | 1.61 | 28.91 | 2.59 |
| 2 | 180% ET | cane | 16.09 | 4.52 | 28.36 | 3.91 |
| 2 | 180% ET | coarse root | 19.06 | 8.58 | 37.33 | 1.75 |
| 2 | 180% ET | fine root | 15.08 | 7.74 | 33.13 | 5.98 |
| 2 | 180% ET | leaf | 51.51 | 3.64 | 3.48 | 1.74 |
| 2 | 180% ET | trunk | 18.13 | 4.33 | 32.52 | 2.44 |
| 3 | 60% ET | arm | 19.09 | 4.10 | 27.78 | 1.51 |
| 3 | 60% ET | cane | 18.02 | 5.11 | 27.70 | 2.06 |
| 3 | 60% ET | coarse root | 18.06 | 3.40 | 38.82 | 3.01 |
| 3 | 60% ET | fine root | 18.07 | 5.20 | 33.50 | 5.04 |
| 3 | 60% ET | leaf | 64.14 | 14.04 | 3.47 | 3.30 |
| 3 | 60% ET | trunk | 20.41 | 5.74 | 29.69 | 2.58 |
| 3 | 120% ET | arm | 19.27 | 2.06 | 31.20 | 3.09 |
| 3 | 120% ET | cane | 19.31 | 5.30 | 28.22 | 1.38 |
| 3 | 120% ET | coarse root | 19.83 | 3.95 | 37.67 | 4.85 |
| 3 | 120% ET | fine root | 19.22 | 4.77 | 39.10 | 3.08 |
| 3 | 120% ET | leaf | 57.39 | 11.98 | 1.53 | 2.39 |
| 3 | 120% ET | trunk | 20.11 | 6.87 | 32.33 | 2.75 |
| 3 | 180% ET | arm | 19.18 | 2.94 | 31.13 | 4.58 |
| 3 | 180% ET | cane | 20.19 | 5.70 | 32.07 | 5.54 |
| 3 | 180% ET | coarse root | 22.09 | 5.90 | 35.02 | 12.11 |
| 3 | 180% ET | fine root | 19.74 | 4.62 | 30.39 | 11.03 |
| 3 | 180% ET | leaf | 66.97 | 16.15 | 3.80 | 3.68 |
| 3 | 180% ET | trunk | 25.90 | 13.65 | 33.76 | 3.62 |
